# Choose your partner: Social evaluation of skilfulness at cooperative co-action tasks in Tonkean macaques (*Macaca tonkeana*)

**DOI:** 10.1101/2024.07.18.604048

**Authors:** Marie Hirel, Hélène Meunier, Roger Mundry, Hannes Rakoczy, Julia Fischer, Stefanie Keupp

## Abstract

Social evaluation – inferring individual characteristics of others from their past behaviours – is an adaptive strategy that helps to inform social decisions. However, how nonhuman primates form and use impressions about others to select their social partners strategically is still unclear. In this study, we investigated whether Tonkean macaques, *Macaca tonkeana*, can spontaneously use information, acquired by observation, to choose the optimal partners for cooperation in two co-action tasks from the same domain. The subjects (N=5) did not prefer to cooperate with the skilful partner compared to the unskilled partner, irrespective of the task or how much attention they paid to the partners’ actions solving a solo version of one task prior to the test. The probability of optimal choices did not increase through trials either, indicating no learning by experience with the partners across the 16 test trials. Our results contradict findings of previous studies that tested monkeys in different domains of competence and contexts, and thus encourage further investigation on monkeys’ social evaluation abilities. Our experimental design presents a promising way to investigate different contexts, the inter-individual differences, the different types of social information involved, and the cognitive mechanisms underlying social evaluation and partner selectivity in nonhuman primates.

## Introduction

Many nonhuman primates live gregarious lifestyles in which they are confronted with social decisions on a daily basis. For example, they choose conspecific partners in agonistic contexts such as recruitments for coalitions [1–5], in affiliative contexts such as grooming [6] as well as for cooperation [7,8], or to learn foraging techniques from [9,10]. Such decisions may be based on existing social bonds and the individual’s knowledge of its group members’ relationships [11–14]. In addition, social decisions may be based on the assessment of other individuals’ characteristics. For example, knowing about physical strength or skills of a conspecific is advantageous for deciding whether to enter a competition over resources. In some situations, such as recruiting a partner for cooperation, picking a model to learn from, or choosing a partner to form coalitions with, individual characteristics may be as important as relationship status for optimal outcomes. Information about others can be obtained from interactions, observations, or, specifically in humans, gossip from third-party individuals [15]. Social evaluations are an important part of human cognitive capacities and inform decisions on whom to trust, whom to work with, whom to learn from, or whom to compare to in current as well as future encounters [16–18]. Yet, whether and to what extent social information is gathered by nonhuman primates and how it affects their partner selectivity is not yet fully understood.

Several nonhuman primates seem to possess social evaluation skills at least to some degree. For example, they differentiated between prosocial/cooperative and antisocial/uncooperative individuals. Chimpanzees (*Pan troglodytes*) preferentially chose “nice” over “mean” experimenters [19–21], brown capuchins (*Sapajus apella*) preferentially avoided non-helpful experimenters [22], and common marmosets (*Callithrix jacchus*) preferred to approach individuals who engaged in positive cooperative vocal interactions [23]. Common marmosets, brown capuchins, and squirrel monkeys (*Saimiri sciuerus*) also discriminated against non-reciprocators by accepting less food from a human who refused to exchange food with a third party before [24–26]. This preference for prosocial and cooperative social agents supports the hypothesis that partner choice based on social evaluation is a key adaptive strategy promoting and maintaining cooperation [27,28]. For successful cooperation, however, skilfulness in the task at hand is also important; consequently, it’s advantageous to evaluate and consider other individuals’ skill sets when recruiting collaborators.

Nonhuman primates indeed seem to form impressions about the expertise of others and target their attention towards these ‘experts’ in non-cooperative contexts. For example, long-tailed macaques (*Macaca fascicularis*), vervet monkeys (*Chlorocebus pygerythrus*), and Guinea baboons (*Papio papio*) changed their behaviour toward the only individual in the group that was able to provide food in experimental situations [29–31]. Long-tailed macaques also differentiated between an experimenter who repeatedly proved capable of opening a box with food rewards and an unsuccessful experimenter who failed at it [32]. Brown capuchins preferentially observed the more skilful nutcrackers within their group, rather than the lesser skilled ones, independently of social affiliations or social proximity [33,34], and chimpanzees preferentially copied the most expert members of their group at a foraging technique [9,10]. Yet, whether nonhuman primates can evaluate the skilfulness of partners to select them strategically for cooperation is still an open question. Only chimpanzees have been tested in this context and they strategically selected a conspecific cooperator who had successfully helped them in the same situation in the past [35,36]. A recent study extended these findings by showing that directly experiencing outcomes of successful interactions was not necessary for chimpanzees to evaluate conspecifics’ skills in cooperative tasks [37].

Despite these findings, the cognitive mechanisms underlying nonhuman primates’ social evaluation skills require further investigation. Particularly, it is still an open question whether primates can infer future behaviour only for similar matching behaviours or whether they can make wider generalisations of a model’s characteristics and resulting behaviours across contexts. From the human literature, we know about at least three inference types that play a role in social decision-making: behaviour matching, *i.e.*, inferences based on the similarity of the past and current situations, global impression formation, *i.e.*, making wider generalisation of a model’s characteristics across contexts, and rational trait reasoning, *i.e.*, rational trait-like inferences in context-specific ways [17,38–40]. For nonhuman primates, it is difficult to disentangle whether they express social reasoning skills corresponding to either of these inferences or rather to associative learning strategies from their past interactions [41]. For instance, the response pattern of chimpanzees in Melis and colleagues’ study [35] spoke in favour of decisions based on success in the previous trial rather than in favour of a general evaluation of skilfulness. Chimpanzees indeed appeared to rely on both direct interactions [35] and indirect third-party observations [19–21]. By prompting chimpanzees to recruit conspecific partners to perform with at different co-action tasks in cooperation as well as competition, Keupp & Herrmann [37] recently provided evidence that chimpanzees can draw domain-specific inferences about conspecifics’ skills.

In the current study, we investigated whether Tonkean macaques (*Macaca tonkeana*) can choose optimal partners in cooperative co-action tasks and whether they spontaneously (*i.e.*, from the first trials) use information acquired by observation to do so. Subjects had several opportunities to sample information about two human partners’ skilfulness (demonstration phase). One partner was a skilful performer whereas the other failed most of the time. Subjects then had to choose one of the partners to cooperate with on a familiar co-action task (subjects observed the partners operating this task) and on a novel one (subjects had never seen partners operating this task before). We were interested in their partner choices - the optimal strategy to maximise reward outcomes being to pick the partner who was formerly skilful. Optimal choices from the first trial on would indicate a spontaneous preference for the skilful partner based solely on observation, suggesting an evaluation of the partners’ skills, while optimal choices increasing with trials would indicate a learning effect based on direct experience with the partners. We also investigated whether the subjects’ attention during the demonstration phase was predictive of the probability to make optimal partner choices. Contrasting subjects’ partner choices in a novel task condition and a familiar task condition allowed us to investigate whether they generalised their social evaluation across different tasks from the same domain.

We studied this question in Tonkean macaques, a species living in multi-male multi-female societies which is characterised as highly tolerant and can display cooperative behaviours [42–44]. In a tolerant social system, individuals can frequently interact with group members, also outside their own matriline, without risking severe aggression [44–46]. Consequently, they can interact with a broad range of individuals in diverse situations, where knowledge about characteristics of these individuals might come in handy. Tonkean macaques have also demonstrated good knowledge of individuals’ social relationships in their group [14] as well as expressed other social cognitive skills [47–49]. In light of the social and cognitive characteristics of this species and previous findings revealing social evaluation abilities in other nonhuman primates, we predicted that our subjects would preferably choose the skilful partner from the very first trial on, in both tasks. We also expected that the more an individual observed the partners during the demonstration phase, the more relevant information this individual can obtain and, therefore, the greater the probability that this individual makes optimal choices afterward.

## Methods

### 1 Subjects and study site

This study was conducted at the Centre de Primatologie – Silabe de l’Université de Strasbourg, in France. All the subjects lived in one social group, living in a wooded outdoor enclosure of 3788 m^2^ with constant access to an indoor room of 20 m^2^. Individuals were fed once per day with dry pellets and once per week with fruits and vegetables, and had access to water *ad libitum*. They participated in cognitive experiments on a voluntary basis. At the start of the training, the group consisted of 31 individuals (17 females), aged from three months to 27 years. We started the training and familiarisation with 14 individuals in experimental rooms situated next to their outdoor enclosure (see Figure S1 in the Supplementary Materials), but only five of them (one female; age range: 10-24 years; Table 1) managed to pass all familiarisation steps to be tested as subjects in the experiment. The tested individuals had participated in several ethological studies before [*e.g.*, 14,49–52], but they had never been tested with social evaluation experiments before.

**Table 1:** Overview of subjects and their choices. Presented are subject characteristics, the identity of the human partners, test condition (familiar or novel), and how often subjects chose the skilful partner during initial preference assessment and test trials.

| Name | Sex | Age (years) | Initial preference (# skilful) | Skilful partner | First task | Test session 1 (# skilful) | Test session 2 (# skilful) |
| --- | --- | --- | --- | --- | --- | --- | --- |
| Abricot | M | 10 | 2/8 | Actor A | Novel | 4/8 | 6/8 |
| Alaryc | M | 10 | 1/8 | Actor A | Familiar | 2/8 | 3/8 |
| Barnabé | M | 9 | 4/8 | Actor B | Novel | 2/8 | 6/8 |
| César | M | 8 | 3/8 | Actor B | Familiar | 3/8 | 4/8 |
| Néréis | F | 24 | 3/8 | Actor B | Novel | 6/8 | 6/8 |

### 2 Materials

#### 2.1 Puzzle box tasks

Subjects were tested with two different types of puzzle boxes: the Slider task and the Cog task (Figures 1a and 1b, and [37]). These tasks required subjects to navigate a ball inside the boxes and to decide where to move the ball to overcome obstacles and traps. The apparatuses had a size of 0.25 m x 0.25 m and were fixed to the mesh of the experimental rooms from the outside. The monkeys could use their fingers to operate the ball through the mesh. A ball of 28 mm diameter could be inserted on top of the boxes. Subjects had to move the ball from top to bottom and were rewarded once the ball fell into an aluminium tray at the bottom.

**Figure 1:**
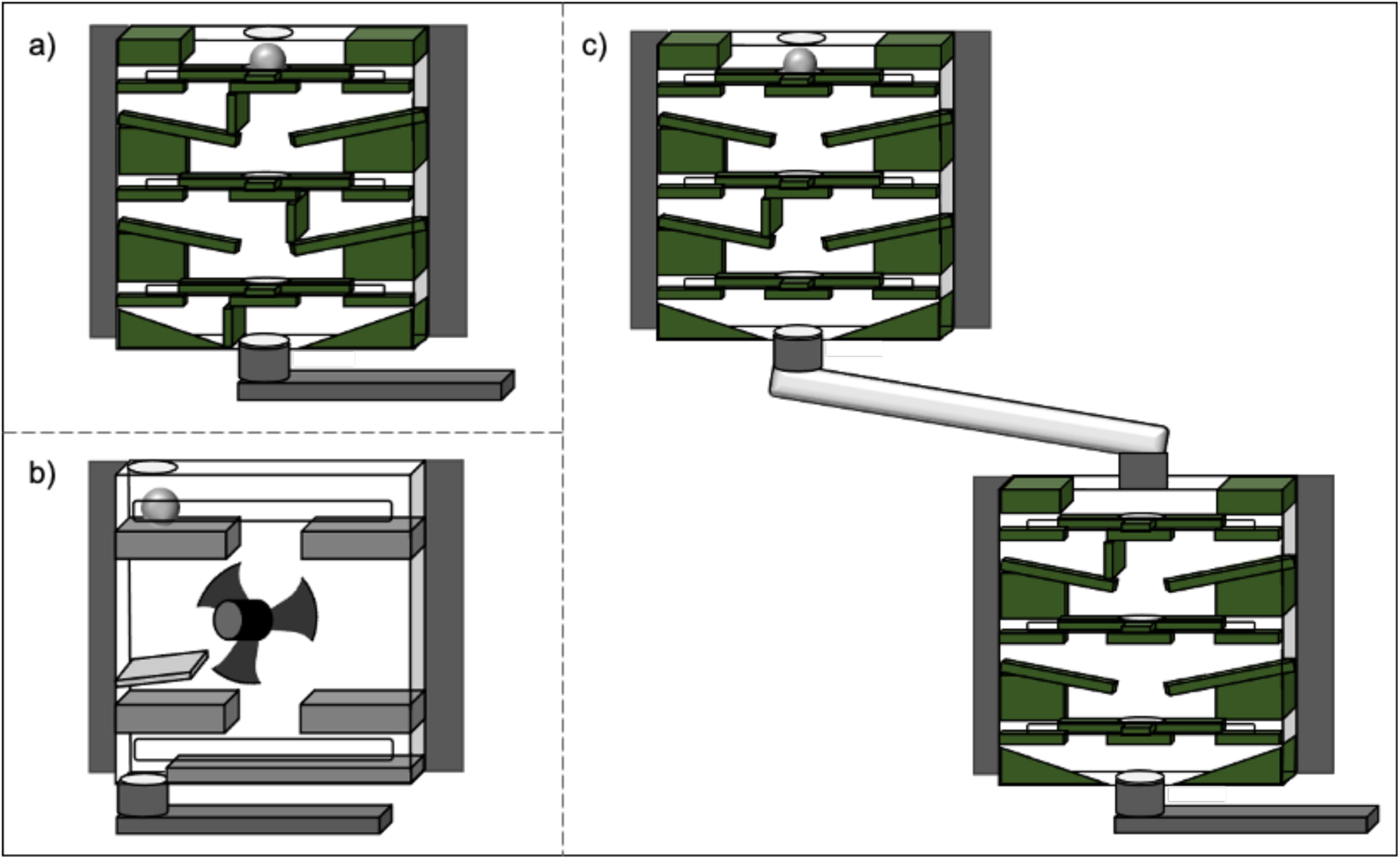
Puzzle box tasks: (a) the single setup of the Slider (familiar) task, (b) the single setup of the Cog (novel) task, and (c) the cooperative co-action setup of the Slider (familiar) task. For the Slider task, traps could be inserted in the box under each level, on either side, to block the ball from falling down to the next level. For the Cog task, a trap could be inserted in the box under either side of the cog wheel to block the ball from falling down.

The puzzle boxes required different actions. The Slider task was a transparent puzzle box with three levels. On each level, the ball was caught on a slider. By moving the slider to the left or to the right, the ball could be transported towards a gap that let it fall down to the next level. Under each level, traps could be inserted on either side to block the ball from falling down to the next one. If the ball was navigated to the blocked side, it was trapped and the attempt was considered a failure. The Cog task required moving the ball from the top level into the middle level, where a cog wheel had to be turned left or right to move the ball further down. Traps could be inserted on either side of the cog wheel: once placed, they blocked the ball from falling down on the respective side and the ball could not be retrieved.

#### 2.2 Cooperative testing setups

A cooperative setup consisted of two puzzle boxes connected by a transparent hose (Figure 1c). One box was mounted slightly higher such that a ball could roll from the upper box into the lower box. If the ball was navigated successfully to the bottom of the upper box, it rolled into the lower box. If the ball was navigated into a trap in the upper box, it never reached the lower box. The two boxes of a cooperative setup were accessible from different positions/rooms, such that each player only had access to one of them and the overall success depended on both players’ successful task performance. During the test phase, two cooperative setups of the same task were mounted on each side of the experimental room (Figure 2). The subjects had to decide with which of two human partners they wanted to perform the task by approaching the respective location and touching the target presented by the human partner.

**Figure 2:**
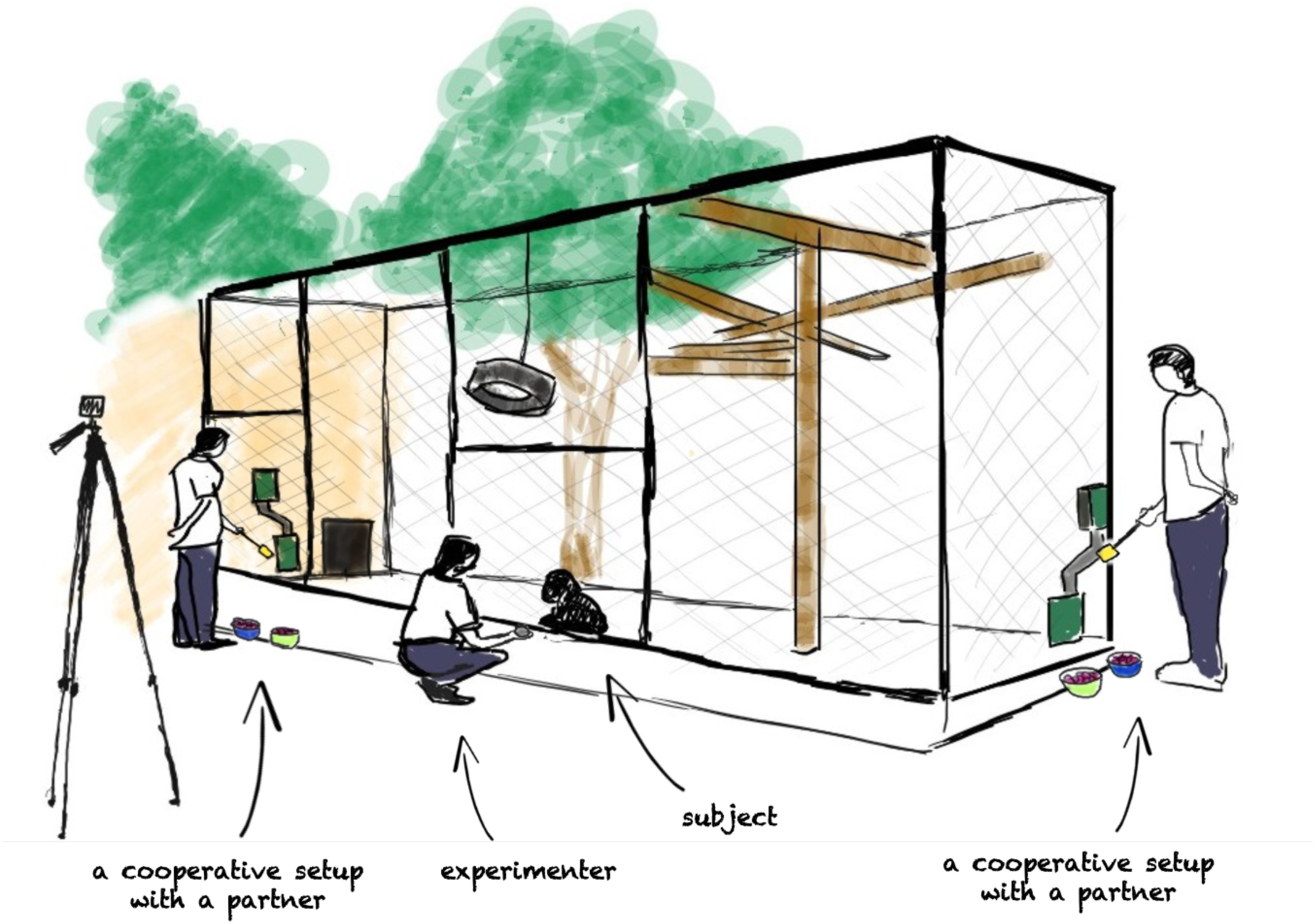
Schematic depiction of the enclosure at the start of a test trial: two cooperative setups were fixed on the left and right side, the experimenter and the subject were situated in the middle equidistant to the two setups, the two partners moved in front of the setups and presented their target (yellow pane). The subject then had to move toward one of the partners and touch the target, indicating its choice to perform the task in cooperation with this partner. Illustration by J. Desriac

### 3 Procedure & design

Subjects underwent an extensive familiarisation phase with comprehension checks to ensure they understood the tasks (for details, see the Supplementary Materials). Specifically, they learned to operate the single setups (*i.e.*, individual box), to navigate the cooperative setup by operating the boxes sequentially, and to indicate their choice by touching a target at the desired location. They also had to demonstrate their principal understanding of the consequences of choosing a cooperative setup with or without a co-action partner in place by choosing the correct setup, i.e., they had to reliably choose the location with a partner who could access the upper box of the cooperative setup rather than the location with no partner in place.

#### 3.1 Initial preference assessment

Two unfamiliar humans played the role of the partners. To assess whether subjects had an initial preference for one of the partners, we conducted eight initial preference trials prior to experimental manipulation. The experimenter lured the subject in the middle of the test room before each trial (to be at an equal distance from the partners). Then, both partners approached the mesh and presented a raisin on the palm of their hand. The subject could approach one of the partners to take their raisin; this approach was considered as a choice for this partner. The partners’ position was randomised across trials, with an equal number of times on each side. If the subjects demonstrated no preference for one partner, the role of the partners was randomly assigned to the subjects and counterbalanced between subjects. If the subject chose one partner five times or more out of the eight trials, we assigned to this ‘preferred’ partner the role of the unskilled partner for this subject. In this way, the subjects had to change their preference toward the other partner to succeed in the cooperative test. As no group’s initial preference for the same person was found, the roles of the partners could be counterbalanced between subjects (Table 1).

#### 3.2 Demonstration phase

Subjects were able to observe the partners individually performing at the Slider task to gather information on the partners’ skills. Subjects gained information about the partners when these were playing alone on single setups of the task – in contrast to the co-action setups used for the cooperative testing condition. Our decision on the amount of exposure to partners’ skills was based on other studies such as Melis and colleagues [36] and Subiaul and colleagues [21], in which chimpanzees required at least twelve trials of interactions with each partner. Since we were specifically interested in whether monkeys spontaneously use information acquired merely by observation to pick a co-action partner, we went beyond this minimum amount and used 24 information sampling events per partner. These trials were divided into three sessions of eight trials, with one session conducted per day. The partners’ presentation order and location were alternated and randomised across sessions (for more details, see section 3 and videos in the Supplementary Materials).

In each demonstration session, the skilful partner was successful in all eight trials whereas the unskilled partner was instructed to fail in seven out of eight trials, e.g., by navigating the ball into a trap. Hence, by the end of the third session, subjects observed the skilful partner succeeding in 24 out of 24 (100 %) trials and the unskilled partner succeeding in three out of 24 (12.5 %) trials. Once the skilful partner successfully navigated the ball through the puzzle box, the experimenter uttered her praise and excitement to emphasise the positive and desired outcome, followed by a provision of a reward (one grape) to the skilful partner. The unskilled partner had 30 s for trying to navigate the ball down without success, followed by the experimenter verbally emphasising the negative outcome.

#### 3.3 Test phase

Subjects had to choose one of the partners to cooperate at the co-action tasks. They were tested for two test sessions: one session with the Slider task (familiar condition) and one session with the Cog task (novel condition). Subjects were tested with both partners. Two cooperative setups of one of the two tasks were installed, one on each side. Each session consisted of eight trials, and we presented one session per day. A trial refers to the choice to work on the co-action task with one of the partners. The presentation order of novel and familiar sessions was counterbalanced across subjects: two subjects were first tested with the familiar task, and three subjects were first tested with the novel task. The partners’ location was randomised across trials, with an equal number of times on each side across subjects.

A trial began with the experimenter luring the subject to a central position equidistant to each cooperative setup. Then, both partners moved simultaneously to their designated setup and presented a target (a long wooden stick with a yellow square at the end). The subjects indicated their choice by approaching their preferred location and touching the respective target. Only then, the experimenter moved in front of the chosen cooperative setup and inserted the ball into the partner’s box. The unchosen partner moved away from the mesh and returned to their initial position. If the skilful partner was chosen, subject and partner successfully navigated the ball through the setup and received a reward (a grape) from the experimenter. The skilful partner received the same reward as the subject to emphasise the cooperativeness of the task. If the unskilled partner was chosen, the partner failed at navigating the ball successfully and the ball never reached the subject’s part of the setup. The experimenter waited for 30 s after the insertion of the ball before ending the trial. In this case, neither the unskilled partner nor the subject were rewarded. The skilful partner was always successful (100 %) during these two test sessions whereas the unskilled partner was never successful (0 %) (see the videos in the Supplementary Materials).

### 4 Data coding

All sessions were videotaped with one to four cameras (GoPro Hero8) depending on the experimental step to obtain different views of the subjects, the partners, setups, and experimental rooms. Two different observers, including one who was unaware of the study design and hypothesis, coded independently frame by frame all the videos of the three experimental phases using Behavioral Observation Research Interactive Software (“Boris”; [53]). Videos were coded for: (a) the partner choices of the subjects at the initial and test sessions (inter-coder reliability: Cohen’s kappa, *κ* = 1, *N* = 118), and (b) the duration of looking behaviours of the subjects (i.e., head orientation, or eye orientation when visible) directed toward a partner (face, body, or hands) during each demonstration trial (inter-coder reliability: *ICC* = 0.893, *N* = 240). During the demonstration phase, only one partner was present at a time. We only coded subjects’ looking time when the partner was performing the task, not when it was the subject’s turn (for more details, see the Supplementary Materials).

### 5 Data analyses

#### 5.1 Did the subjects prefer the skilful partner over the unskilled partner?

We aimed to investigate whether Tonkean macaques could spontaneously use information acquired by observation to choose optimal partners to cooperate with at a familiar and a novel co-action task. A spontaneous preference for the skilful partner would manifest in optimal choices at the first trial and no effect of sessions or trials on subjects’ choices. A transfer of social evaluation of partners’ skills across the two different tasks would manifest in no effect of task on subjects’ choices. However, estimating the effects of task (novel, familiar), session (1, 2), trials, and their potential interaction on the probability of choices of the skilful partner led to a model too complex for our small final sample size. We hence revised our analysis plan to investigate whether the subjects chose the skilful partner more often in both test sessions, regardless of the task, and how their choices developed across trials (*i.e.*, with increasing direct experience with the actors). We fitted a model in R (version 4.3.2; [54]) using Generalized Linear Mixed Models (GLMM; [55]). We used the function glmer of the package lme4 (version 1.1-35.1; [56]) and a binomial error structure with logit link function [57]. We estimated the effect of session (test session 1, test session 2) on the probability to choose the skilful partner. To control for their potential effects, we included into this model subjects’ initial partner preference and trial number as fixed effects, with trial number in an interaction with session.

The reason for including the interaction between session and trial number was that we expected the effect of learning through trials to be more pronounced in the first session than in the second. For instance, while only having information on the partners’ skills from observations when starting the first session, subjects then obtained additional information from direct interactions with the partners at each trial of the first session that they could use for the next trial. Since the same subjects were tested several times in both sessions and in order to avoid overconfident model estimates and to keep type I error rate at the nominal level of 5 %, we included subject ID as a random intercepts effect and all identifiable random slopes [58,59], namely those of trial number and session. As an overall test of the fixed effects and to avoid cryptic multiple testing [60], we conducted full-null model comparisons using likelihood ratio tests [61]. We compared this full model with a null model lacking the effect of session and its interaction with trial number in the fixed effects part but being otherwise identical. We tested the effect of individual fixed effects by comparing the full model with a reduced model lacking the interaction in the fixed effects’ part [59]. We obtained confidence intervals of model estimates and fitted values by means of a parametric bootstrap (N=1,000 bootstraps; function bootMer of the package lme4). We checked all the relevant model assumptions and transformed some variables when needed to ease the interpretation of the model estimates and convergence (for more details, see section 5 in the Supplementary Materials).

#### 5.2 Was the attention of the subjects predictive of their partner choices?

We fitted a similar model except that we estimated the effect of attention during the demonstration on the number of choices of the skilful partner in the test phase. To potentially obtain the relevant information on the partners’ skills, subjects needed to observe both partners during the demonstration phase. As a measure of the subjects’ attention to both partners, we hence chose the minimum looking time per subject to the skilful and unskilled partner, *i.e.*, the time during which each subject observed each partner, and used this for the following analysis. Into the model, we included the attention during the demonstration as a fixed effect, test session (test session 1, test session 2), and their interaction. The reason for including the interaction between attention and session was that we expected the effect of learning through sessions to be less pronounced for subjects with a higher level of attention during the demonstration sessions compared to less attentive subjects, as attentive subjects could already know at the first test session which partner is the optimal choice and then do not have to learn from direct experience. We again included subject ID as a random intercepts effect and test session as a random slope within subject. We compared this full model with a null model lacking the effect of proportion of total attention and its interaction with session in the fixed effects part but being otherwise identical and with a reduced model lacking the interaction (for more details, see section 6 in the Supplementary Materials).

## Results

### 1. Did the subjects prefer the skilful partner over the unskilled partner?

Subjects chose the skilful partner 17 out of 40 times (42.5 %) in Test session 1 and 25 out of 40 times (62.5 %) in Test session 2 (Table 1). Overall, their probability to choose the skilful partner was not significantly influenced by session and its interaction with trial number (full-null model comparison: *χ^i^*= 4.55, *df* = 2, *p* = 0.103; Tables 2 and 3), indicating that our subjects’ probability to choose the skilful partner did not change over the course of trials and sessions (*i.e.*, with the acquisition of direct experience with the partners). Although all subjects chose the skilful partner more often in the second test session, this increase was marginally non-significant (*p* = 0.069; Table 3, Figure 3 and Figure S2 in the Supplementary Materials). In addition, the probability to choose the skilful partner at the very first trial was not different from chance (Figure 3). However, one subject stood out in choosing the skilful partner in six out of the eight trials in both test sessions (*i.e.*, familiar and novel tasks), and beginning from the first trial. This subject was presented with the novel task at her first test session. She also successfully reversed her actors’ preference from the initial preference session in which she had chosen the human who would play the skilful actor three out of eight times (Table 1). On a group level, the strength of the subjects’ initial preference for one actor did not significantly influence their probability to make optimal choices after (Table 3).

**Figure 3:**
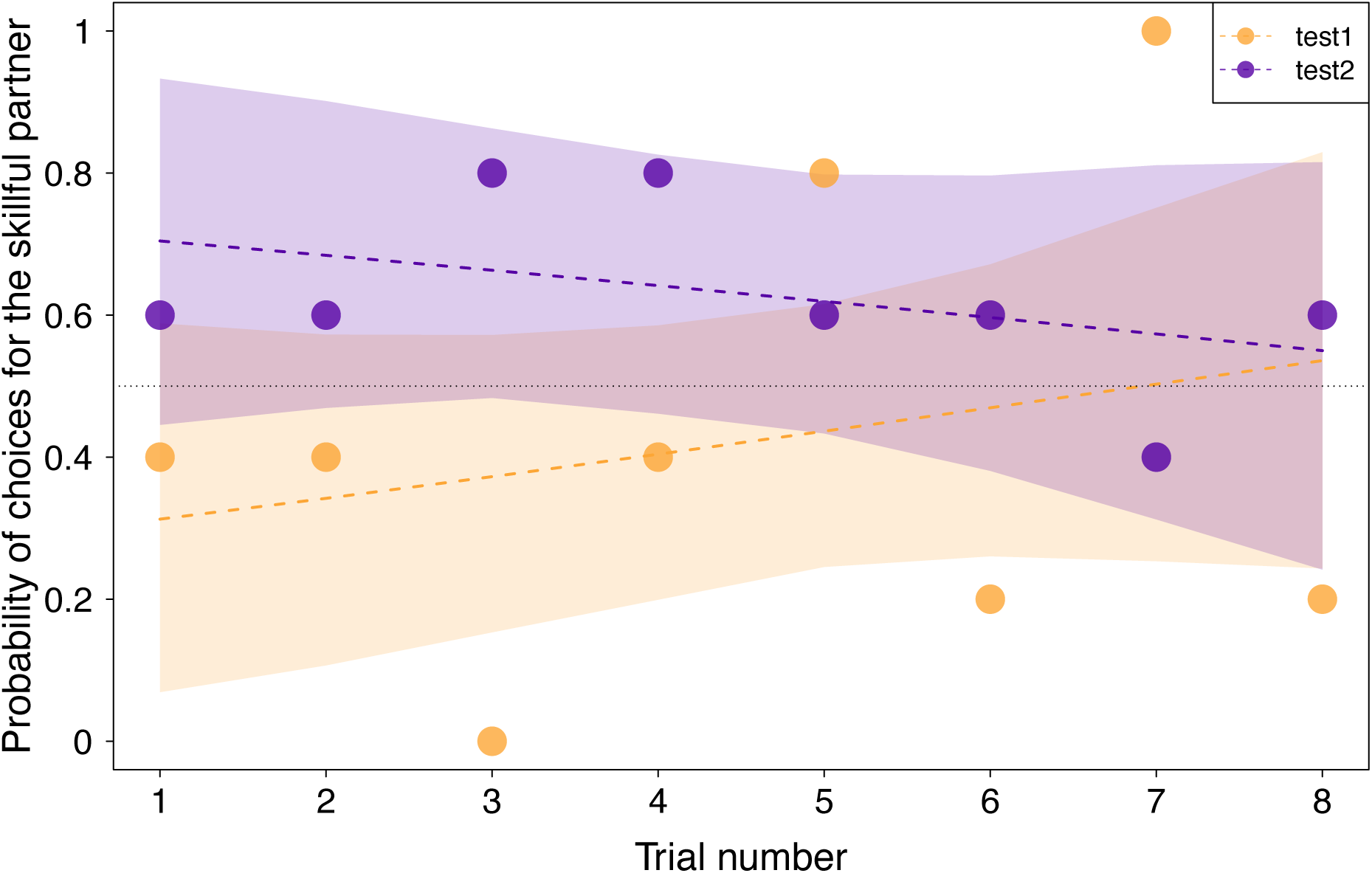
Probabilities of subjects’ choices for the skilful partner. The raw data and fitted model and its 95 % confidence limits are shown for both sessions for initial preference being at its average. Each data point represents the mean probability of choosing the skilful partner for the five subjects at each trial. Note that at the first trial, the confidence intervals of both sessions encompass the value of 0.5 indicating that the performance in the first trial did not significantly deviate from chance.

**Table 2:**
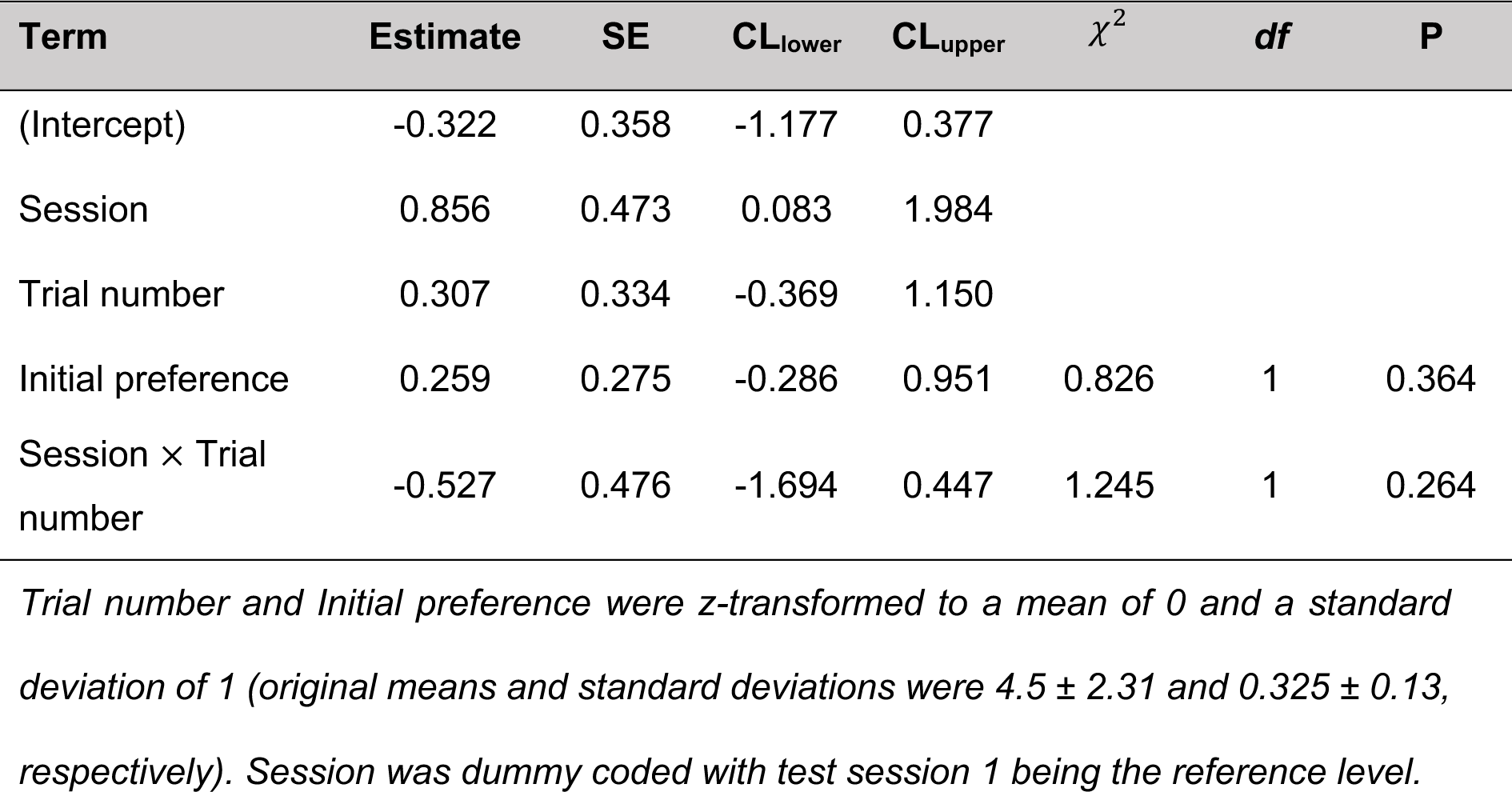
Results of the subjects’ partner choices full model (estimates together with standard errors, 95 % confidence limits, significance tests, and the estimates’ range after excluding individuals one at a time).

| Term | Estimate | SE | CL<sub>lower</sub> | CL<sub>upper</sub> | $\chi^2$ | df | P |
| --- | --- | --- | --- | --- | --- | --- | --- |
| (Intercept) | -0.322 | 0.358 | -1.177 | 0.377 |  |  |  |
| Session | 0.856 | 0.473 | 0.083 | 1.984 |  |  |  |
| Trial number | 0.307 | 0.334 | -0.369 | 1.150 |  |  |  |
| Initial preference | 0.259 | 0.275 | -0.286 | 0.951 | 0.826 | 1 | 0.364 |
| Session × Trial number | -0.527 | 0.476 | -1.694 | 0.447 | 1.245 | 1 | 0.264 |

*Trial number and Initial preference were z-transformed to a mean of 0 and a standard deviation of 1 (original means and standard deviations were $4.5 \pm 2.31$ and $0.325 \pm 0.13$ , respectively). Session was dummy coded with test session 1 being the reference level.*

**Table 3:** Results of the subjects’ partner choices, reduced model (estimates together with standard errors, 95 % confidence limits, and significance tests).

| Term | Estimate | SE | CL<sub>lower</sub> | CL<sub>upper</sub> | $\chi^2$ | df | P |
| --- | --- | --- | --- | --- | --- | --- | --- |
| (Intercept) | -0.315 | 0.352 | -1.166 | 0.437 |  |  |  |
| Session | 0.843 | 0.468 | 0.103 | 2.018 | 3.304 | 1 | 0.069 |
| Trial number | 0.048 | 0.234 | -0.455 | 0.577 | 0.041 | 1 | 0.839 |
| Initial preference | 0.254 | 0.270 | -0.339 | 0.848 | 0.826 | 1 | 0.363 |

*Trial number and Initial preference were z-transformed to a mean of 0 and a standard deviation of 1 (original means and standard deviations were $4.5 \pm 2.31$ and $0.325 \pm 0.13$ , respectively). Session was dummy coded with test session 1 being the reference level.*

### 2. Was the attention of the subjects predictive of their partner choices?

During the demonstration phase, subjects looked on average 6.1 ± 2.9 s out of 16.3 s per demonstration trial at the skilful partner and 11.1 ± 6.9 s out of 26.7 s at the unskilled partner. In total, they looked on average 145.4 s at the skilful partner (range across individuals: 125.5 s to 182.9 s), 265.5 s at the unskilled partner (range across individuals: 209.9 s to 370.4 s), and 410.9 s at both partners (range across individuals: 335.4 s to 505 s) across all demonstration trials. Overall, the probability to choose the skilful partner was not significantly influenced by attention and its interaction with session (*χ^i^* = 2.123, *df* = 2, *p* = 0.346; Tables S3 and S4 in the Supplementary Materials), indicating that neither the attention of the subjects to both partners during the demonstration phase nor its interaction with test session influenced their partner choices. Therefore, contrary to our expectation, the individuals who observed the partners more during the demonstration did not make more optimal partner choices in the test sessions.

## Discussion

We investigated whether Tonkean macaques can choose optimal partners for cooperation in two co-action tasks. Subjects had several opportunities to observe a skilful and an unskilled partner performing individually at a solo version of a task (familiar task) and then had to choose one of them to cooperate with in the familiar task and a novel task from the same domain. During each test trial, the subjects obtained information on the partners’ skills through direct experience – receiving food only if they chose the skilful partner. Optimal choices from the first trials would then indicate a spontaneous preference for the skilful partner based solely on observation, suggesting an evaluation of the partners’ skills. Optimal choices increasing with trials would indicate a learning effect based on direct experience with the partners. Contrasting subjects’ partner choices in a novel task condition and a familiar task condition allowed to investigate whether they generalised their social evaluation across different tasks from the same domain.

We found that Tonkean macaques did not prefer to cooperate with a skilful human partner compared to an unskilled partner. Overall, they did not spontaneously use information acquired by observation prior to the test to choose optimal partners, indicating no evaluation or use of knowledge on the partners’ skills. Even though they tended to choose the skilful partner more often in their second test session compared to the first session, this increase in optimal choices across sessions was marginally non-significant. This result indicates no learning by experience with the partners across the eight or 16 presented test trials. Our subjects’ choices emerged irrespective of how much attention they paid to the partners’ actions when they observed the partners solving solo versions of the task in the demonstration phase. Contrary to our predictions, Tonkean macaques who observed the partners the most during the demonstration then did not make more optimal choices thereafter.

These findings must be carefully interpreted as resulting from a small sample size and with noticeable inter-individual differences in our subjects’ partner choices. Three subjects chose the skilful partner six out of eight times only in the second test session and two subjects did not make more than four optimal choices in both sessions. One individual stood out by choosing the skilful partner above chance at both test sessions and from the first trial on. This choice pattern could indicate that she transferred her knowledge from the observation phase to the test phase and the two co-action tasks, but it could also be the result of chance. However, this Tonkean macaque chose again six out of eight times a skilful over an unskilled actor at opening containers in another experiment (Hirel *et al.*, in prep.). This inter-individual variation in performance should be investigated in the future by testing the same individuals in different settings. Our paradigm presents a promising way to address this question, as being suitable to test individuals with different partners pairs (humans as well as conspecifics) differing in their degree of competence or social relationship and with different tasks and in different contexts, like competition (see [37]).

In addition, our subjects were attentive to the partners’ actions less than 50 % of the time. This lack of interest in partner performance is in accordance with previous findings showing that long-tailed macaques only paid close attention to partner performance if it had directly relevant consequences for themselves [62,63]. Indeed, in our study, the partners’ performance during the demonstration phase did not affect the subjects’ food intake, which might have led to a lack of sufficient interest in our subjects. However, we had no predictions on how much time Tonkean macaques should observe the partners’ actions as we still do not know how many seconds of demonstrations primates need to observe to obtain the relevant information about others. This question would have to be assessed in the future with a different study design that systematically varies the amount of information provided.

The overall lack of optimal partner selectivity in our experiment stands in contrast with previous findings with monkeys [22,24–26]. However, contrary to our experiment, these previous studies presented a larger amount of test trials, each including an observation of the actors’ actions and, directly after, a choice to take a food piece from one of the actors. For example, capuchins and squirrel monkeys had to choose between a reciprocator and a non-reciprocator in twelve sessions of twelve trials each [24,25]. In addition, their subjects’ choices had no effect on their food intake as they always got the same piece of food no matter which actor they chose. Therefore, these monkeys may not have used associative learning despite the large trial number, but they had 144 opportunities to observe each actor compared to 24 possible observations in our study. Except for one study indicating no change in capuchins’ choices against non-helpers through sessions [22], information on how the choices of these monkeys developed through trials is not provided in the respective papers. Knowing the development of these monkeys’ choice pattern would help us understand whether we did not offer sufficient opportunities for Tonkean macaques to acquire relevant knowledge about the partners’ skills, or whether other reasons may explain our contrasting findings.

One possible reason for the differences in monkeys’ social evaluative performance between our experiment and previous studies may also be the investigation of different domains of competence and contexts. Indeed, these previous studies tested monkeys on prosocial behaviours which relate to a different domain than foraging or cooperative skilfulness and may require different cognitive capacities. The monkeys in previous studies could observe human experimenters interacting with each other prosocially or not and then decide who they will accept food from. In contrast, Tonkean macaques in our experiment observed the partners individually performing the task and then decided which partner they wanted to cooperate with. Taking food from an actor’s hand might be a less cognitively challenging situation than having to choose a partner for a cooperative activity. Indeed, choosing a cooperative partner can be complex as it may involve taking into account not a single but a broad range of social information as well as interacting and coordinating with another individual [17].

Another possibility is that skilfulness might not be a salient feature at all to assess others’ value for cooperative tasks for monkeys as for apes or humans. To our knowledge, evidence of active cooperative foraging in wild monkey species populations has never been described. However, monkeys pay attention to the foraging behaviours of others and adapt their own behaviours toward the foraging experts of their group in non-cooperative contexts [29–31,33]. Skilfulness of others might then be a relevant characteristic for monkeys to consider in contexts other than cooperation. For example, being able to evaluate individuals’ skills and use it to selectively learn new behaviours from the best experts can be truly advantageous. However, social learning in primates has mainly been studied with a focus on the learning mechanisms (what to learn) or the social aspects (who to learn from), but only taking into account the social relationships between individuals and never the individual characteristics [64–66]. Future research could investigate whether monkeys form and use impressions of others’ expertise to choose from whom to learn new behaviours and how this selective learning through social evaluation might affect the evolutionary dynamics of innovation and culture.

In summary, we investigated Tonkean macaques’ ability to choose optimal partners in a cooperative context using evaluation of partners’ skilfulness acquired by observation. Even though our subjects did not prefer to cooperate with a skilful partner, showing no evidence of social evaluation abilities, the inter-individual differences on our small sample size encourage to further investigate whether and how Tonkean macaques can form impressions about others on different behaviours and contexts. Besides, our methodology represents a promising way to investigate different contexts, the different types of social information involved, and the cognitive mechanisms underlying social evaluation and partner selectivity in nonhuman primates.

## Acknowledgments

We are thankful to the management and staff of the Centre de Primatologie – Silabe de l’Université de Strasbourg (https://www.silabe.com) for their authorisation, support, and help with the data collection, especially the valuable assistance of the animal keepers, the technical team, and the researchers and students. A very special thanks to Pascal Ancé and Pierre-Henri Moreau for agreeing to play the roles of the partners in the experiment. For their invaluable support with the conception and the manufacturing of the experimental apparatuses, we would like to thank Louis Frank and M. Zippert and his team of the chemistry workshop of the University of Göttingen. We are grateful to Adam Provin, Michele Marziliano, and Rose Riguet for their precious help with the data collection, and Nadja Vöglte for reliability coding. Finally, we thank Julie Desriac for the illustration of the experimental setup.

## Ethical statement

This study respects the European ethical standards and regulations (Directive 2010/63/UE) and has been approved by the internal ethical committee of the Centre de Primatologie – Silabe de l’Université de Strasbourg (SBEA 2022-03, registration n° B6732636). We only used positive reinforcement and all individuals participated on a voluntary basis, *i.e.*, they were never forced to enter, be isolated, or to perform the tasks in the experimental rooms. Their daily feeding regime was not affected by cognitive testing and water was available *ad libitum* at all times.

## Data accessibility

The supplementary document, videos, data, statistical code associated with this article can be found at: https://osf.io/tnkz2/ (DOI: 10.17605/OSF.IO/TNKZ2).

## Authors Contributions

**Hirel Marie**: Conceptualization, Data curation, Formal analysis, Investigation, Methodology, Project administration, Visualisation, Writing – original draft, Writing – review & editing. **Meunier Hélène**: Resources, Writing – review & editing. **Roger Mundry**: Formal analysis, Writing – review & editing. **Rakoczy Hannes**: Conceptualization, Funding acquisition, Writing – review & editing. **Fischer Julia**: Funding acquisition, Writing – review & editing. **Keupp Stefanie**: Conceptualization, Methodology, Project administration, Supervision, Writing – review & editing.

